# Meningeal Worm Infection in Central Iowa Goat Herds II: Individual Cases and Treatment Using a Camelid Therapeutic Protocol

**DOI:** 10.1101/613562

**Authors:** Joe Smith, Amanda Kreuder, Ryan Breuer, Kelly Still-Brooks

## Abstract

Meningeal worm (*Paralaphostrongylus tenuis*) infection, also known as cerebrospinal nematodiasis, is a common parasitic infection in New World Camelids in the United States. There is also a considerable risk for this disease in the Boer goat population. Despite the rapidly increasing size of the national goat herd, there are no treatment protocols reported in the literature for goats with this disease. This study describes a successful clinical approach and treatment of 3 Boer goat cases with therapy previously reported for use in New World Camelids. The clinical presentation, diagnosis, and long-term outcome of *P. tenuis* infections in these goats presented to ISU Food Animal and Camelid Hospital (FACH) is reported here within. Practitioners should be aware that clinical presentation and diagnosis are similar for goats as reported for camelids with cerebrospinal nematodiasis. Additionally, the described treatment protocols for camelids appear to demonstrate a comparative efficacy in goats.

## Introduction

Complete cases, including treatment and outcome of cerebrospinal nematodiasis caused by *Parelophostrongylus tenuis* (more commonly known as “meningeal worm” infections) in goats are lacking, with few reported in the literature. Those cases that are reported in the peer-reviewed literature focus on pathological findings, with a paucity of descriptions of neurological progression, treatment success or clinical outcome. Currently there are limited cases of successful treatment of cerebrospinal nematodiasis in goats reported, with none utilizing the current therapies used for llamas and alpacas, and as such clinicians are limited to comparative literature when attempting treatment for such cases.

Infection with *P. tenuis* occurs naturally in White Tailed Deer (WTD) in the eastern and Midwestern United States. The parasite is host adapted such that infected deer often display no clinical signs, and experience extremely low morbidity and mortality in deer. The problem arises when hosts other than WTD are infected, as these other hosts often experience significant morbidity and mortality from the migrating *P. tenuis* larvae.

At the present time, no anti-mortem “gold standard” test for the diagnosis of cerebrospinal nematodiasis exists. The presence of larvae in the central nervous system upon gross necropsy, or the presence of larval genetic material (via PCR) in central nervous tissue of a deceased animal remain the definitive diagnostic standard. This presents a challenge when live animals are presented with clinical signs suggestive of cerebrospinal nematodiasis. Fortunately for veterinary medicine, the presence of clinical signs such as ataxia and recumbency, coupled with an eosinophilic pleocytosis, in an area that is endemic to *P. tenuis* is recognizable as the standard for ante-mortem diagnosis.

The recent arrival of South American Camelid species, such as the llama and the alpaca, in the US have highlighted the risks of *P. tenuis* infection for animals that serve as aberrant hosts for this parasite. A recent treatment protocol has shown promise for the treatment of Camelids with cerebrospinal nematodiasis. The purpose of this study was to exam the effect of the camelid treatment protocol on goats with clinical signs of cerebrospinal nematodiasis.

### South American Camelid Treatment Protocol

Treatment of cerebrospinal nematodiasis in South American Camelids is described with three principles of therapy^1^: Anti-inflammatory drugs; anthelmintics; as well as exercise restriction/physical therapy.

#### Anti-inflammatories

The non-steroidal anti-inflammatory drugs flunixin meglumine (1.1-2.2 mg/kg/day, IV)^7^, and meloxicam (1 mg/kg/day, SC)^2^ have been reported for use in camelids. Steroidal anti-inflammatories such as dexamethasone have also been reported, although they have been linked to an increased mortality rate.^1^

#### Anthelmintics

Ivermectin (0.2 mg/kg, once SC) and fenbendazole (50 mg/kg/day x 5 days, PO)^7^ have been utilized to kill any larvae still aberrantly migrating through the host in camelids.

#### Exercise Restriction and Physical Therapy

Due to the neurologic dysfunction, exercise restriction in the form of separation from the herd is recommended. Also, due to this compromise, passive range of motion and stretching is utilized to avoid complications from recumbency in camelids.

## Materials and Methods

The medical records of cases admitted to the Food Animal and Camelid Hospital (FACH) of Iowa State University Lloyd Veterinary Medical Center (ISU LVMC), were generated for all goats over a period of time from January 1 2000 through January 1 2018. These records were scoured for diagnosis of diseases of the nervous system, consistent with cerebrospinal nematodiasis. Cases were confirmed with either cerebrospinal eosinophilia in the patient, or in a similarly presenting herd mate on a farm where cerebrospinal nematodiasis had been recently been diagnosed. Incomplete records that did not fulfill the admission criteria were removed from consideration of the study.

Information on signalment, history, physical examination findings, diagnostic tests, clinical diagnosis, treatment, and response to treatment were gathered from medical records used in this study. Clients were consulted for follow-up information regarding the case post discharge.

## Results and Discussion

### History, Physical Examination and Diagnostics

Cases: 3 goats were identified that met the admission criteria for this retrospective case series.

Case 1: A 9 month old intact male Boer buck presented to the ISU Lloyd VMC FACH for evaluation following a progressive lameness and ataxia of approximately 72 hours duration. The buck presented with bilateral hind limb paresis in a “dog-sitting” posture. Neurological examination yielded absent proprioception of the hind limbs, decreased pelvic muscle tone, decreased patellar reflexes, as well as tachycardia (presumptive from handling and examination). Other exam findings were within normal limits. The buck was from a herd of 40 goats with no other immediate medical concerns, and had been purchased three months prior. This case’s CSF tap is demonstrated in Table 1.

**Table 1:**
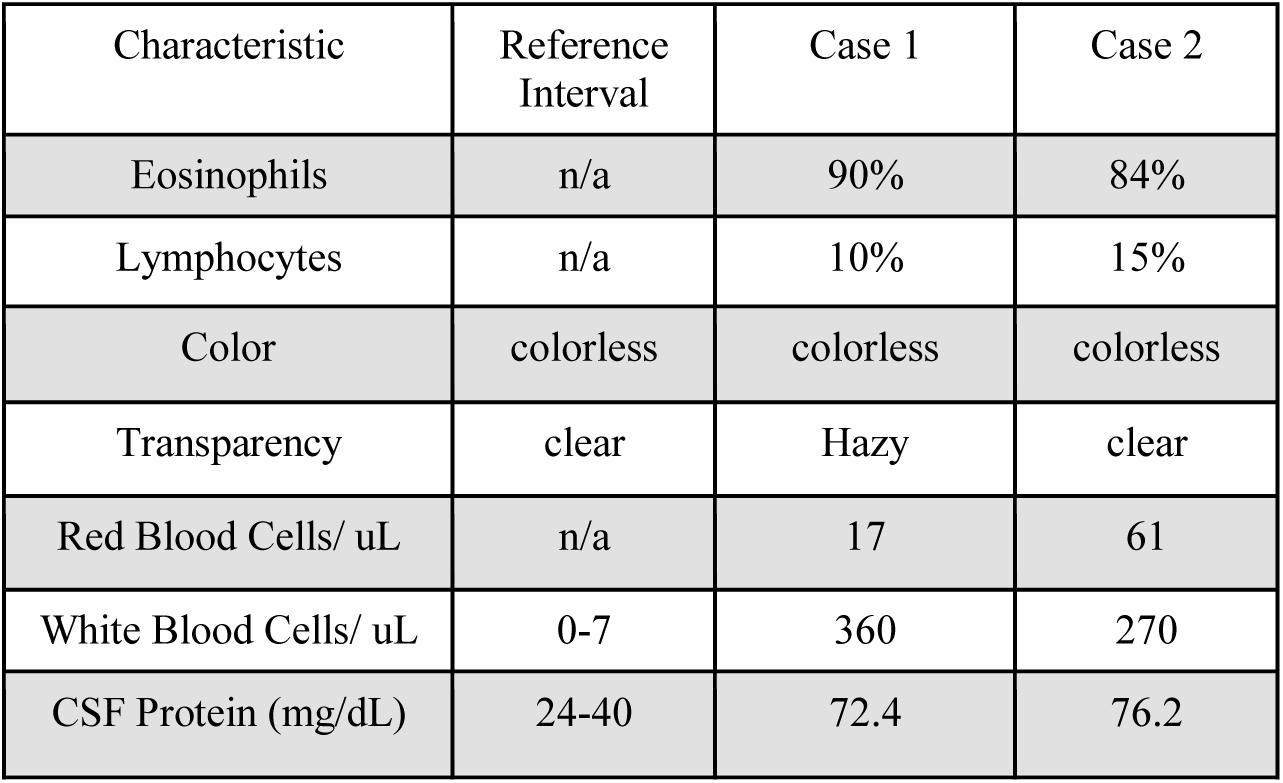
Cerebrospinal Fluid Characteristics of Case 1 and Case 2

| Characteristic | Reference Interval | Case 1 | Case 2 |
| --- | --- | --- | --- |
| Eosinophils | n/a | 90% | 84% |
| Lymphocytes | n/a | 10% | 15% |
| Color | colorless | colorless | colorless |
| Transparency | clear | Hazy | clear |
| Red Blood Cells/ uL | n/a | 17 | 61 |
| White Blood Cells/ uL | 0-7 | 360 | 270 |
| CSF Protein (mg/dL) | 24-40 | 72.4 | 76.2 |

Cases 2 & 3: Presented at the same time as members of the same herd. They are members of a herd of 13 animals that share a low-lying pasture that floods in the spring. Other members of the herd have had neurologic signs prior including weakness, ataxia, cranial nerve deficiencies and opisthotonus.

Case 2: Case 2 was a 6 month old female Boer goat who presented to ISU FACH for a 4 day history of hind limb lameness. On physical examination she had difficulties with conscious proprioception and coordination of her thoracic and pelvic limbs, although this was more pronounced in the pelvic limbs. Her physical examination was otherwise unremarkable. Her CSF is demonstrated in Table 1.

Case 3: This case was a 7 month old intact male Boer goat that presented to the ISU FACH for signs of ataxia and a head tilt over a 14 day period. A CSF tap on this patient was declined due to financial reasons, but due to the proximity and temporal association with Case #2 a presumptive diagnosis was made. Note: while other neurological diseases, such as Listeriosis, could cause a presentation such as the head tilt seen in Case #3, this animal did not demonstrate other clinical signs of listeria (circling, nystagmus) and responded to a treatment for P. tenuis infection, which would not be expected to demonstrate efficacy for a bacterial infection such as listeriosis.

Cerebrospinal Fluid (CSF) Collection and Analysis: Prior to CSF collection patients were sedated with midazolam (0.5 mg/kg, IV) or a combination of midazolam (0.5 mg/kg, IV) and butorphanol (0.05 mg/kg). The lumbosacral space was clipped, and after an initial skin prep a 1-2 mL infiltration of 2% lidocaine was subcutaneously infiltrated. The site was then sterile prepped via alternating scrubs of chlorhexidine and 70% isopropyl alcohol. The skin incised with a stab incision of a #15 blade. A spinal needle was then introduced and directed until CSF fluid was observed in the hub of the spinal needle. Serial samples were then collected and saved in commercial plasma and EDTA tubes. The samples were then analyzed for cell count via a cytospin preparation and a cell count by a board-certified veterinary clinical pathologist. The CSF was also analyzed for protein count, cellularity, and color.

### Therapeutic Results

All 3 goats tolerated treatment well. None of the commonly associated signs of toxicity for the reported drugs [neurotoxicity for ivermectin, and abomasal (gastric) ulceration for meloxicam] were appreciated. Additionally all three goats tolerated physical therapy exercises well.

All cases were prescribed meloxicam (1 mg/kg PO q 24-48 hours) for extended periods of time after discharge. This extended duration was reported by the clients to also be efficacious and safe with no evidence of abomasal ulceration (melena, bruxism) observed while returned to the farm. All three goats dramatically improved over the initial hospitalization period and all three survived to discharge. The improvement of neurologic parameters over the hospitalization of Case 1 is presented in Table 2.

**Table 2:** A selected example of the progression of neurologic signs for Case 1 during the first 5 days of therapy.

|  |  |  |  |  |  |
| --- | --- | --- | --- | --- | --- |
| Patellar Reflexes<br>(LPL: Left Pelvic Limb; RPL: Right Pelvic Limb) | LPL – Decrease response but present;<br>RPL – Decrease response but present | LPL – Weak, decrease response but present;<br>RPL – Weak, decrease response but present | LPL – Very weak, decrease response but present;<br>RPL – Improved reflex response in right hind limb, still decreased response | LPL – Present;<br>RPL – Present | LPL – Present;<br>RPL – Present |
| Conscious proprioception<br>(hind limbs) | Absent conscious proprioception in hind limbs | Absent conscious proprioception in hind limbs | Absent conscious proprioception in hind limbs | Absent conscious proprioception in right hind limb but present; weak proprioception in left hind limb | Conscious proprioception present in the hind limbs with slight delay |
| Perineal Reflexes | Anal tone decreased, normal perianal sensation present, and adequate tail tone | Anal tone decreased slightly, normal perianal sensation present, and adequate tail tone | Anal tone normal, normal perianal sensation present, and adequate tail tone | Anal tone normal, normal perianal sensation present, and adequate tail tone | Anal tone normal, normal perianal sensation present, and adequate tail tone |

Case 1: 21 days post hospitalization case 1 demonstrated improved ambulation, but still had evidence of ataxia as well as an inability to stand for extended durations of time. 3 months past discharge from the hospital no ataxia was present and at 5 months post discharge he returned to his normal status in the herd.

Cases 2 and 3: Both cases improved over the duration of their hospitalization. Case 2 returned to normal ambulation within 3 months of discharge from the hospital. Case 3 returned to normal ambulation within 2 months of discharge.

Preventive strategies as described for camelids (monthly subcutaneous injections of ivermectin) were discussed with the clients, but ultimately not performed as resistance to anthelmintics is a serious concern for small ruminants and there is currently no data to support the use of this practice in species other than camelids.

Both clients instead utilized preventive strategies of minimizing exposure to the arthropod vector by limiting access to standing water, moving pastures frequently, fencing to minimize deer contact and vigilant observation of the rest of the herd.

No additional cases from either herd have been reported. Within 2 years of these cases being presented to the FACH.

## Conclusions

We observed multiple similarities in the clinical signs of cerebrospinal nematodiasis in goats as well as the utility of a camelid-based treatment protocol for the therapy of this condition in goats. Cerebrospinal nematodiasis in goats appears to clinically present similarly to camelids. Signalment, history, neurological exam and cerebrospinal fluid analysis are key to antemortem diagnosis.

Treatment as described for llamas and alpacas, (ivermectin, fenbendazole, flunixin meglumine, and meloxicam) was attributed with a successful outcome. It is noteworthy to mention that this therapy involves usage of drugs and doses not labelled for goats, therefore strict accordance to AMDUCA with respect to extra-label drug use is warranted.As described in other species, full neurological improvement, if it will occur, may take several months’ time.

